# Genetic, Genomic and Biophysical Case Study: Familial 46, XY Sex Reversal due to a Novel Inherited Mutation in Human Testis-Determining Factor SRY

**DOI:** 10.1101/2021.05.05.442859

**Authors:** Elisa Vaiani, Yen-Shan Chen, Pablo Ramirez, Joseph Racca, Maria Sonia Baquedano, Carmen Malosetti, Maria Sol Touzon, Roxana Marino, Mariana Costanzo, Marcela Bailez, Esperanza Berensztein, Maria Laura Galluzzo-Mutti, Deepak Chatterjee, Yanwu Yang, Alicia Belgorosky, Michael A. Weiss

**Author notes:** Corresponding Authors: Alicia Belgorosky, MD, PhD, ORCID ID: orcid.org/0000-0002-4234-400X, Michael Weiss, MD, PhD, ORCID ID: orcid.org/0000-0003-1325-1610, Research Career Award, Superior Investigator National Research Council of Argentina, Associate Scientific Director, Research Unit Garrahan-CONICET, Ex Chair Woman of Endocrinology Department, Director and Professor of Pediatric Endocrinology Career, School of Medicine, University of Buenos Aires (UBA), Argentina. Contributed equally as co-first authors (Elisa Vaiani, Yen-Shan Chen). Contributed equally as co-senior authors (Alicia Belgorosky, Michael A. Weiss). Disclosure summary: The authors declare that they have nothing to disclose.

## Abstract

**Objective:** To describe the clinical, histopathological and molecular features of a novel inherited *SRY* allele (p.Met64Val; consensus box position 9) observed within an extensive pedigree: two 46, XY sisters with primary amenorrhea (16 and 14 years of age; probands P1 and P2), their normal father and brother, and an affected paternal XY grandaunt.

**Design, Setting, Participants and Outcome Measurements:** Following DNA sequencing to identify the SRY mutation, hormonal studies of the probands and histopathological examination of their gonads were performed. Functional consequences of p.Met64Val (and other mutations at this site) were also investigated.

**Results:** Breast development in P1 and P2 was Tanner II and IV, respectively. Müllerian structures and gonads resembling ovaries were found in each sister. Histopathology revealed gonadal dysgenesis, gonadablastoma and dysgerminoma. AMH/MIS, P450 SCC, and P450 aromatase were expressed in gonadoblastoma tissues. Genomic sequencing revealed no candidate mutations in other genes related to sexual differentiation. Variant p.Met64Val impaired *Sox9* transcriptional activation associated with attenuated occupancy of the testis-specific enhancers *Enh13* and *TESCO*. Biophysical studies indicated that the mutant HMG box retains specific DNA binding and DNA bending but with accelerated rate of protein-DNA dissociation.

**Conclusion:** The partial biological activity of p.Met64Val SRY, maintained at the threshold of SRY function, rationalizes opposing paternal and proband phenotypes (the “the father-daughter paradox”). Steroidal biosynthesis by gonadoblastoma may delay genetic diagnosis and recognition of gonadal tumors. Quantitative assessment of inherited SRY alleles highlights the tenuous transcriptional threshold of developmental decision-making in the bipotential gonadal ridge.

## Introduction

The male phenotype in therian mammals is determined by *SRY*, a small, single-exon gene located in the <u>s</u>ex-determining region of the <u>Y</u> chromosome ^1; 2^. SRY is expressed in the embryonic gonadal ridge just prior to its morphologic differentiation to initiate a sequence of regulatory steps that simultaneously promotes testis development and suppresses ovarian development in the early bipotential gonad ^3^. Subsequently, anti-Müllerian hormone/Müllerian inhibiting substance (AMH/MIS) and testosterone (T) respectively secreted by Sertoli cells (AMH/MIS) and Leydig cells (T) in the fetal testis promote regression of Müllerian primordia and external male differentiation ^4^.

SRY is an architectural transcription factor (TF) and member of the SOX family that contains a central high-mobility group (HMG) box. This conserved motif ^5^, containing three α-helices in an L-shaped structure, binds within an expanded DNA minor groove to introduce sharp bends ^6^. Such DNA binding and bending are sequence specific. The transport of SRY into the nucleus is directed by two distinct nuclear localization signals (NLS), located at either end of the HMG-box domain ^6; 7^.

The monobasic C-terminal NLS bindsβ1 importin (imp-β1NLS; ^8^) whereas the N-terminal NLS binds exportin-4 ^9; 10^ and calmodulin ^11; 12^.The HMG box of human SRY also contains a nuclear export signal recognized by exportin-1 (CRM1). Clinical mutations in SRY associated with gonadal dysgenesis have been identified that impair its binding as cargo to imp-β1, exportin-4 and exportin-1 ^6; 13^. Such clinical correlations have established that the developmental function of SRY requires nucleo-cytoplasmic trafficking ^10^, which is coupled to phosphorylation as an activating post-translational modification ^10^. Although the SRY HMG box has also been proposed to mediate protein-protein interactions within the nucleus ^3^, it is presently unclear which candidate partner proteins participate in male-specific gene regulation.

Complementary studies of patients with 46, XY gonadal dysgenesis and related mouse models have provided evidence that SRY directly activates *SOX9*/*Sox9* at the transcriptional level and that such activation is central to testicular differentiation ^14; 15^. Indeed, critical SRY-responsive DNA enhancer elements have recently been defined upstream of the *Sox9*/*SOX9* transcriptional start sites ^14–17^; these are conserved between mouse and human genes. Chromatin-immunoprecipitation (ChIP) assays have shown that SRY binds to specific DNA sites within these control elements (the ancillary testis-specific core element (*TESCO*; ^14^) and critical enhancer element 13 (*Enh13*; ^16^). Loss-of-function mutations in SRY or deletions in *Enh13* lead to failure of testicular differentiation in embryogenesis ^16^. A female somatic phenotype often ensues due to lack of AMH/MIS secretion from the absent Sertoli cells (leading to retention of the Müllerian primordia) and lack of T secretion from the absent fetal Leydig cells (leading to absent genital virilization). Although initial ovarian morphogenesis may occur (as observed in mouse models), on post-natal assessment human patients typically exhibit intra-abdominal streak gonads ^4; 18^.

Genetic screening of patients with differences of sexual differentiation (DSD) have suggested that 10-20% of XY sex-reversal (Swyer Syndrome with complete gonadal dysgenesis; CGD) are due to mutations in *SRY* ^18; 19^. The classical Swyer phenotype consists of bilateral CGD with female external genitalia, uterus, and fallopian tubes ^20^. Partial gonadal dysgenesis is less common and may be associated with formation of ovotestes ^21^. To date, >100 *SRY* variants have been reported, including point mutations, frameshifts and deletions ^22^. The majority arise *de novo (*as meiotic errors in paternal spermatogenesis) and cluster in the HMG-box domain, hence affecting only one individual in a family ^23^. To our knowledge, however, 20 inherited cases have also been reported, creating a seeming paradox (*i.e*., a variant Y chromosome shared by a fertile father and his dysgenetic XY daughter). Although some such cases (n=8) can be explained by paternal mosaicism (the fathers contain both a wild-type [WT] and variant *SRY* ^24–30^, in 9 unrelated families mosaicism was excluded ^31–41^. Molecular mechanisms underlying such variable phenotypes within a pedigree include partial impairment of nuclear import ^42^ and enhanced susceptibility of an unstable SRY variant to proteasomal degradation ^43^, but the majority of such pedigrees have not been investigated at the molecular level. Three additional pedigrees have been reported, but were not well characterized at the clinical or molecular levels ^44–46^.

In the present study we describe variable phenotypes associated with a novel inherited mutation in the SRY HMG box. An integrated analysis of this mutation’s effects on gonadal histology, hormonal pattern and cell-biological properties is presented. The mutation (p.Met64Val; position 9 in the HMG-box consensus sequence ^47^) occurs within the “hydrophobic wedge” motif mediating sharp DNA bending ^48; 49^. Remarkably, the variant *SRY* allele was found in five XY individuals within this pedigree: two 46, XY sisters, their normal 46, XY father and brother, and an affected paternal grandaunt. Whole-genome sequencing of the probands and father excluded DSD-associated mutations in other genes (two neutral polymorphisms were found in one *SOX9* allele in each subject). To investigate how this mutation might perturb gene regulation, we undertook biophysical and mammalian cell-line-based studies in relation to previously described *de novo* clinical variants at the same site in SRY (p.Met64Ile ^50^ and p.Met64Arg ^51; 52^). Whereas the mutation introduced only subtle biophysical perturbations in the DNA-binding properties of the variant HMG box, impaired SRY-dependent transcriptional activation of principal target gene *Sox9* was observed in each case, to an extent (mild or marked) proportionate attenuated occupancy of the testis-specific *Enh13* and *TESCO* enhancer elements. Quantitative assessment of inherited SRY alleles associated with the father-daughter paradox highlights the tenuous transcriptional threshold of developmental decision-making in the bipotential gonadal ridge.

## Materials and Methods

### Patients

This study was approved by the Independent Ethic Committee of the Hospital de Pediatría Garrahan. Informed consent was obtained from the patients and family members.

#### Patient 1

(P1), the index case (proband) was a 16.5-year-old (y) female, referred due to primary amenorrhea. She was born full term (39 weeks) after an uneventful pregnancy. She was the first daughter of healthy unrelated parents. Birth weight was 3.25 kg, and she was discharged after 3 days.

P1 had a feminized phenotype without sexual ambiguity. Perineal inspection revealed normally developed labia majora, minora, and clitoris with normal opening of the urethra and vagina; no dysmorphic features were found. Breast development was Tanner stage II, and pubic hair was Tanner stage III. Her bone age was 14 y. P1 weighted 64 kg (+1.37 SDS), and her height was 161 cm (+0.04, and −1.1 SDS for normal female and male Argentinean references, respectively). Pelvic ultrasound revealed a small uterus (3.9 x 2 x 0.8 cm) and hypoplastic gonads; the left gonad volume was 1.2 ml with multi-cystic internal features whereas the right gonad volume was 1 ml.

#### Patient 2

(P2), the proband’s younger sister (14.3 y) was tested despite signs of normal female puberty. She was born after 40 weeks of an uncomplicated pregnancy. Birth weight was 3.14 kg, and she was discharged after 4 days.

The bone age of P2 was 14 y. She also exhibited a fully developed feminized phenotype without ambiguity. Breast- and pubic hair development were both Tanner stage IV; vaginal mucosa exhibited characteristic features of estrogen stimulation. Menarche was absent. She weighed 40 kg (+1.48S SDS), and her height was149 cm (−1.37 and −3.05 SDS for normal female and male Argentinean references, respectively). P2 had no dysmorphic features. Pelvic ultrasound revealed a pubertal uterus (5.8 x 3.3 x 2.2 cm, endometrial 7.6 mm). The left gonad was enlarged with complex internal cystic features; the right gonad was 1.2 ml.

Sisters, P1 and P2 each had 46, XY karyotype as evaluated in peripheral blood lymphocytes. Levels of serum hormones are given in **Table 1**. In brief, serum LH and FSH levels were high in both patients, suggesting gonadal dysfunction. Serum estradiol levels were undetectable in P1 but within the normal pubertal range in P2. Serum testosterone and inhibin B were undetectable, indicating an absence of a functional testis; AMH/MIS was in each case within the normal female range. Tumor markers in each sister’s serum (β-subunit of human chorionic gonadotrophin (β-hCG) and α-fetoprotein (AFP]) were negative.

**Table 1.** Serum hormone evaluation of the probands at diagnosis

|  | **P1** | **P2** | **RV**<br>**(female)** |
| --- | --- | --- | --- |
| **Chronological age**<br>**(years)** | 16.5 | 14.3 |  |
| **FSH** (mUI/ml) | **61.6** | **73** | 5.88 ± 2.91 |
| **LH** (mUI/ml) | **22.6** | **27,6** | 4.04 ± 3.19 |
| **Estradiol** ( pmol/l ) | ND | 164.8 | 91.7-1505 |
| **Testosterone**( nmol/l ) | ND | ND | 0.17-1.38 |
| **DHEAS**( nmol/l ) | 6697 | 1578.8 | 767-11326 |
| **Cortisol** ( nmol/l ) | 463.5 | 275.9 | 138-607 |
| **17OHP**( nmol/l ) | 1.81 | 3,00 | 1.9-6.1 |
| **Δ4androstenedione**<br>(nmol/l ) | 4.2 | 2 | 1.92-6.9 |
| **Progesterone** (nmol/l) | 0.89 | 0.79 | 0.25-1.2 |
| **βHCG** (mUI/ml) | <5 | <5 | <5 |
| **AFP** (ng/ml) | <3 | <3 | <3 |
| **AMH** (pmol/l) | 30.7 | 21.6 | 5-60 |
| **Inhibin B** (pg/ml) | ND | ND | ND-341 |

P: patient
βHCG: βhuman chorionic gonadotropin
AFP: α-fetoprotein
AMH: anti-Müllerian Hormone
RV: Reference values according to sex and breast Tanner stage of pubertal development
In bold: abnormal results
ND: not detectable

Patients underwent laparoscopic surgery due to the risk of gonadal germ-cell tumors. The right gonad of P1 contained a bilobulated tumor. The right tube and gonad were removed. The left exhibited a streak morphology and was also removed. Müllerian structures were confirmed. No evidence of macroscopic metastases was observed. The left gonad of P2 contained a large, round encapsulated tumor whereas the right gonad contained a tumor with brain-like appearance. Bilateral salpigo-gonadectomy was done. No evidence of macroscopic metastases was observed. Hystopathological evaluation of gonads tissues revealed the presence of gonadal tumors in both P1 and P2 (see Results).

Following surgery, serum estradiol and AMH/MIS levels were undetectable in both patients. Hormone-replacement therapy with estrogen and progestin was subsequently initiated, and menses occurred in each patient. Although no postoperative chemotherapy was indicated, serial measurements of tumor markers (β-hCG and AFP) and pelvic ultrasound examinations were performed to exclude recurrence of gonadal tumors.

An extensive family history was obtained. The ethnic background of the family was Spanish without indication of consanguinity. The first generation was from the north of Spain; the second generation migrated to Argentina. As is shown in **Fig. 1**, three members of the third generation were notable: (i) subject 7 was an 46, XY female with primary amenorrhea and sterility who underwent bilateral gonadectomy at age 24 y; no histopathologic report was available; (ii) subject 1 was a putative XY female without offspring; and (iii) subject 4 was an ostensible male who needed testosterone replacement therapy.

**Figure 1.**
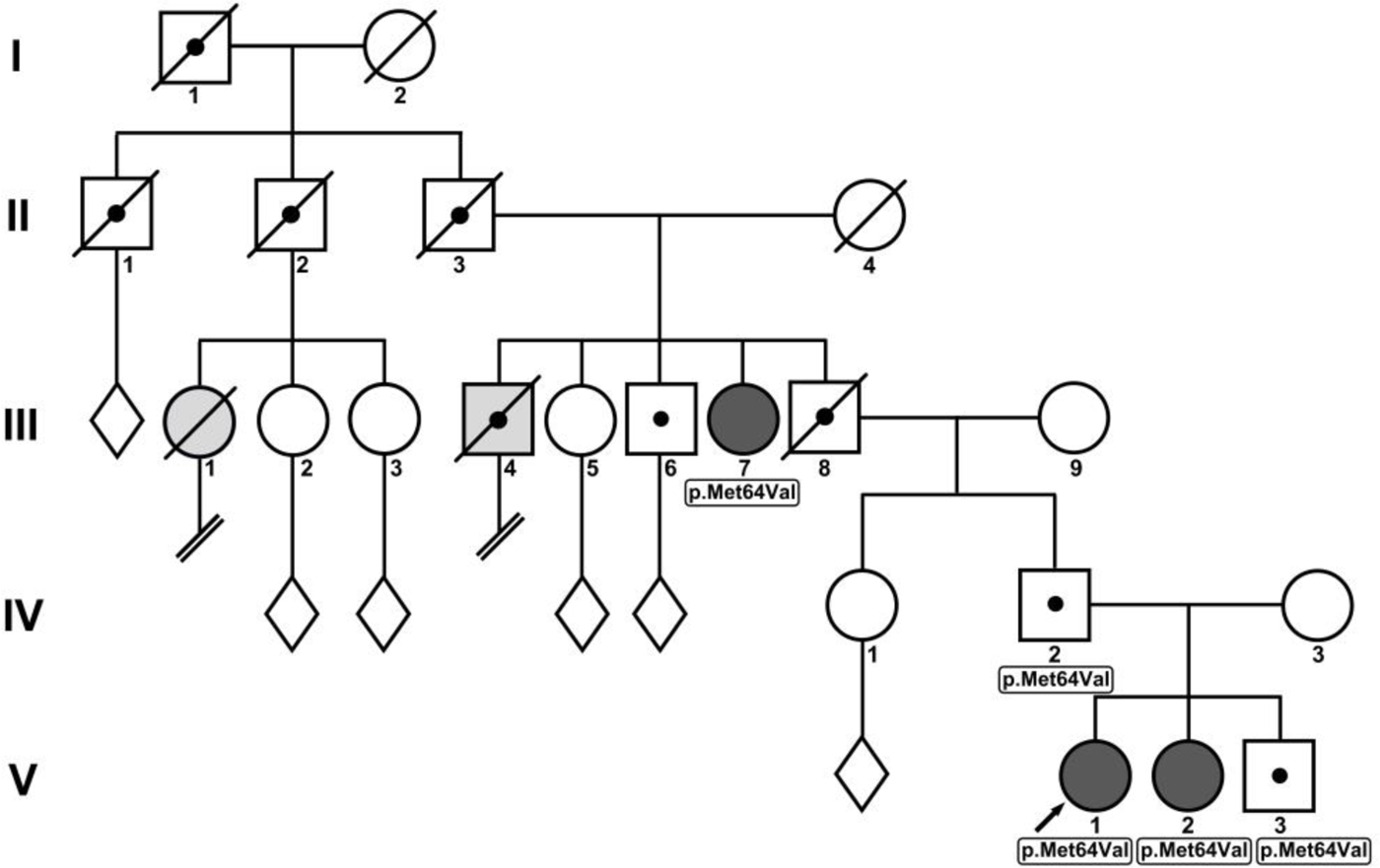
Dark gray circle: Affected proband; Black dot: Obligate carrier with normal male phenotype; White diamond: Unspecified progeny; Double forward slash: Without offspring; Light gray circle: Suspected female phenotype without offspring (E+P substitution); Light gray square: Suspected male phenotype without offspring (T substitution)

### Laboratory analysis

Serum hormone levels were measured as follows: LH and FSH by automated chemiluminescent microparticle immunoassay (ARCHITECT i4000; Abbott); testosterone, estradiol, progesterone, DHEAS, and cortisol by chemiluminescent assay (IMMULITE2000; Siemens); delta 4 androstenedione, 17OH progesterone by RIA (Dia Source) Progesterone; inhibin B and AMH/MIS by 2-site ELISA (Beckman Coulter and Beckman Coulter Gen II, respectively); and βHCG and AFP CMIA-ARCHTECT I 4000.The assays were performed as recommended by the manufacturers. Reference values referred by the manufacture for pediatric population were confirmed in the clinical laboratory of the Hospital de Pediatría Garrahan (Argentina). Bone age was evaluated by the Grewlych and Pyle method.

### Genetic and genomic analysis

Karyotype was determined using peripheral blood lymphocytes; 50 metaphase cells were examined by conventional and G-banding techniques. Genomic DNA from the two sisters, their father, brother, and paternal grandaunt were analyzed. In the first step the coding region of SRY (spanning 204 codons) was amplified by PCR using specific primers as described ^38^. PCR products were purified (QIAquick Gel Extraction Kit, Qiagen, Buenos Aires, Argentina) and sequenced with an ABI PRISM 3500 Genetic Analyzer (Applied Biosystems, Buenos Aires, Argentina) using the Big Dye Terminator v1.1 Cycle Sequencing Kit (Applied Biosystems, Buenos Aires, Argentina). Primers used for sequencing were the same as those used for PCR. The sequences were compared with the NCBI SRY entry (NG_011751.1).

In the second step complete genome sequences of P1, P2 and their father were obtained by next-generation DNA sequencing (based on blood samples) at the Medical Genomics Core at the Indiana University School of Medicine. Genomic DNA was purified by the DNeasy Blood and Tissue Kit Quick-start protocol (QIAGEN). Purified genomic DNA was reconstituted in low-TE buffer. In brief, reconstituted genomic DNA was sheared into small fragments and then ligated to adapters. After hybridization on a flow cell, the fragments were amplified; the sequencer (Illumina model NovaSeq 6000) added labeled nucleotides sequentially and recorded the sequencing reads. The resulting data sets were processed with standard bioinformatics methods following alignment by the Illumina DRAGEN (Dynamic Read Analysis for GENomics) platform. DSD-associated gene sequences (Table 2) were extracted and sorted by specific position on chromosome. The DNA sequences were analyzed by DNA-blast program (National Center for Biotechnology Information; NCBI) to identify the genetic variations, including variation with exonic or non-exonic regions with attention to synonymous and non-synonymous single nucleotide polymorphisms (SNP).

**Table 2.**
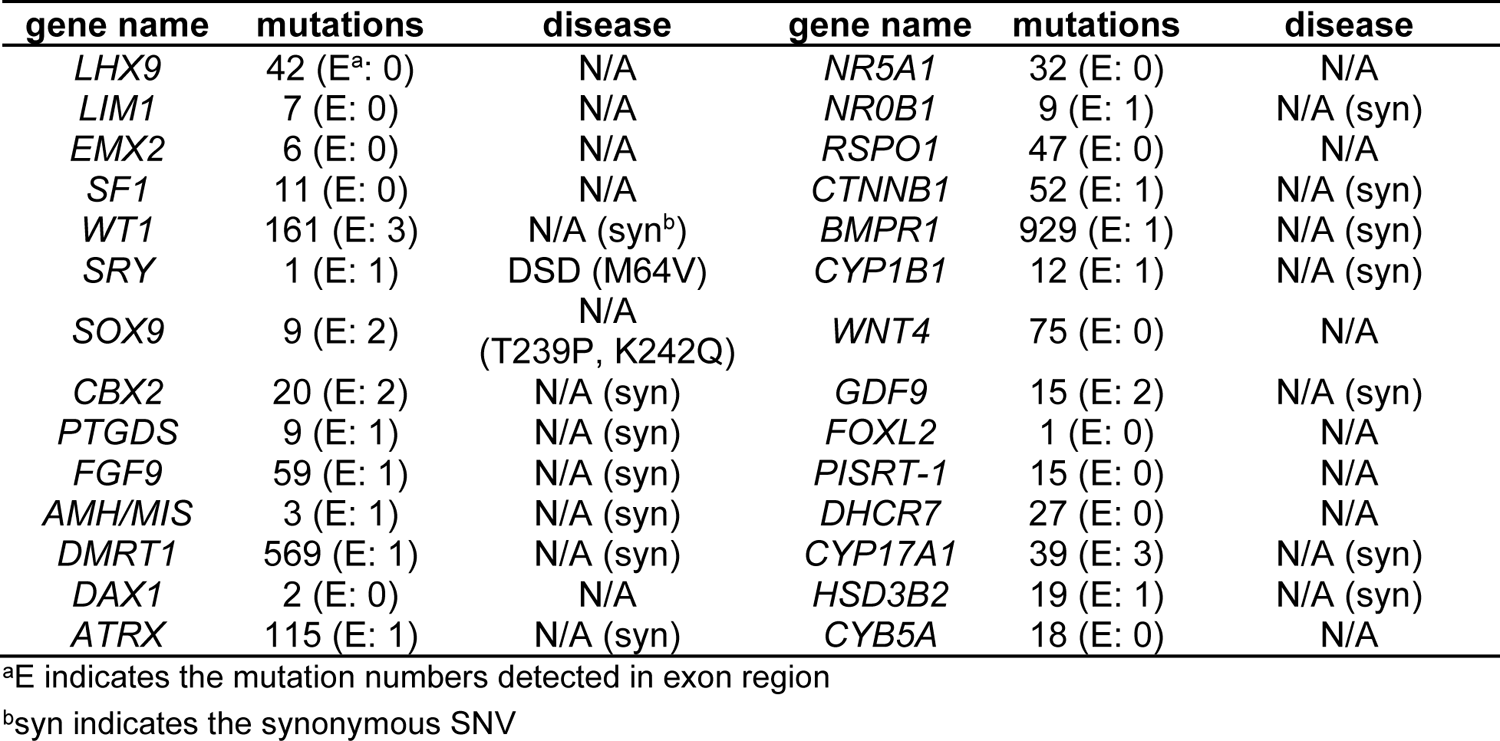
Genomic Analysis of Sex-Related Genes.

### Gonadal histopathology

Gonadal samples were fixed in 10% formalin and embedded in paraffin. Hematoxylin-eosin (H&E) staining and periodic-acid Schiff (PAS) staining were each performed on sections from all surgical specimens. Overall gonadal tissue organization and abnormal histological features were assessed by two specialists in gonadal histology

### Immunohistochemistry (IHC)

PLAP and Cd117 immunostaining were used to characterize gonadal tumors. Pluripotential germ cells were identified using immunohistochemistry (IHC) with anti-OCT3/4 monoclonal antibody (sc-5279, Santa Cruz Biotechnology). Steroidogenic cells were stained with CYP11A1 (P450scc) Rabbit Ab (ab75497, Abcam) (kindly provided by Dr.M. Warman). Estrogen-producing cells were stained by a murine monoclonal antibody recognizing human CYP19 aromatase (ab34193, Abcam) (Dr. M. Warman). Sertoli cells were identified with rabbit Ab AMH (L40) (Dr.Rey R). Germ cells were identified by a murine monoclonal antibody recognizing human placental alkaline phosphatase (PLAP) (Novo Laboratory), and Monoclonal Mouse Anti-Human CD117, conjugated with allophycocyanin, (DAKO Laboratory) Immunohistochemistry (IHC) was performed as described ^53^. In brief, after heat-induced antigen retrieval with a citrate buffer (pH 6.0), sections were incubated overnight with the correspondent antibodies. Staining was performed employing the streptavidin-biotin and peroxidase method, according to the manufacturer’s protocol (DAKO Catalyzed Signal Amplification System, K1500, HRP). After each incubation step, slides were washed in a 0.01% TBS-Tween 20 solution. To provide negative controls, normal serum was used instead of the solution containing a primary antibody; no specific immunoreactivity was detected in these sections. Immunostaining studies were each performed twice, and no differences between duplicates were observed.

### SOX9 Luciferase Assay

Human embryonic kidney cell, HEK 293T, were cultured in Dulbecco’s Modified Eagle Medium (DMEM), supplemented with 10% fetal bovine serum (FBS), 1% penicillin/streptomycin as recommended by the American Type Culture Collection. Cells were transfected with 100 ng of luciferase reporter vector per 1 million cells. The reporter was constructed with 4 of 48-bps elements, derived from the enhancer region of *Col2a1* gene sites. Tested SOX9 variants were co-transfected with 100 ng of plasmid concentration. Normalization control, Renilla plasmid, was also co-transfected in 5 ng of plasmid concentration. Transfections were performed using Lipofectamine 3000 as described by the vendor (Invitrogen). After 8 h in Opti-MEM medium, cells were recovered using fresh culture medium with FBS. Following transient transfection, HEK 293T cells were subjected to the reporter assay. Experiments were conducted in triplicate using the Dual-Luciferase Reporter Assay System (Promega); lysates were simultaneously analyzed for firefly luciferase activity and Renilla luciferase activity (Promega).

### Mammalian plasmids

Plasmids expressing full-length SRY and variants were constructed by PCR and verified by DNA sequencing. Following the initiator Met, the cloning site encoded a hemagglutinin (HA) tag in triplicate to enable Western blotting (WB) and chromatin immunoprecipitation (ChIP).

### Cell culture

Rodent CH34 cells (kindly provided by Dr. P.K. Donahoe, Massachusetts General Hospital; ^4; 10^) were cultured in Dulbecco’s modified Eagle’s medium (DMEM) containing 5% fetal bovine serum (FBS) at 37 °C under 5% CO_2_.

### Transient transfections

Transfections were performed using Lipofectamine 3000 as described by the vendor (Invitrogen). After 8 hours (h) in Opti-MEM medium, cells were recovered using fresh culture medium with FBS. Transfection efficiencies were determined by ratio of green-fluorescent protein (GFP) positive cells to untransfected cells following co-transfection with pCMX-SRY and pCMX-GFP in equal amounts ^10^. Subcellular localization was visualized by immunostaining 24-h post transfection following treatment with 0.01% trypsin (Invitrogen) and plating on 12-mm cover slips. SRY expression was monitored by WB via its triplicate HA tag.

### Chromatin immunoprecipitation

Cells were transfected with epitope-tagged WT or variant SRY. SRY-expressing cells were cross-linked in wells by formaldehyde, collected, and lysed after quenching the cross-linking reaction. Lysates were sonicated to generate proper fragments and immunoprecipitated with anti-HA antiserum (Sigma) containing a Protein A slurry (Cell Signaling). A non-specific antiserum (control IgG; Santa Cruz) served as non-specific control. PCR and qPCR protocols were as described ^43^. Quantification was investigated by qPCR and presentative gel images were collected by Gel Doc (Bio-Rad).

### Cellular fractionation

Cells were treated in the suspension buffer as described ^10^. Lysates were kept on ice, sheared by five passages through a 25-gauge needle, and centrifuged at 2,500 × g for 15 min at 4 °C; supernatants provided cytosolic extract. Pellets were suspended in nuclear lysis buffer, sheared by needle passage, kept on ice for 15 min, and subjected to 13,000 × g centrifugation for 15 min at 4 °C.

### Western blot

24-h post transient transfection, cells were split evenly into 6-well plates and treated with cycloheximide to a final concentration of 20 μg/ml in DMEM for the indicated times; cells were then lysed by RIPA buffer (Cell Signaling Technology). Protein concentrations were measured by BSA assay (Thermo); cell lysates were subjected to 4-20% SDS-PAGE and WB using anti-HA antiserum (Sigma-Aldrich) at a dilution ratio of 1:5000; α-tubulin antiserum provided a loading control. For phosphorylation analysis, HA-tagged SRY variants were immunoprecipitated with rabbit polyclonal anti-phosphoserine antiserum (Abcam). WB following 4-20% sodium dodecyl sulfate-polyacrylamide gel electrophoresis (SDS-PAGE) employed HRP-conjugated anti-HA antibody (Roche). Quantification was performed by Image J software.

### Transcriptional activation assay

In the transient transfections (above) the expression plasmid encoding HA-tagged WT SRY or an HA-tagged variant SRY was diluted 1:50 with the parent empty plasmid to reduce protein expression to the physiological range (*ca*. 10^3^-10^4^ protein molecules/cell ^10^); similar expression levels were verified by anti-HA Western blot. Following transient transfection, cellular RNA was extracted using RNeasy as described by the vendor (Qiagen).SRY-mediated transcriptional activation of *SOX9/Sox9* was measured in triplicate by qPCR as described ^10; 54^. Primer sequences for all of the tested genes were applied as described ^55^. *Tbp*, encoding the specific 5’-TATAADNA-binding subunit of TFIID, was used as an internal control; measurements were made in triplicate.

### Steady-State Fluorescence Spectroscopy

Stability of the free domain was determined using fluorescence spectroscopy and monitoring the Trp emission wavelength 390 nm. Free protein domains were made 5 μM in Buffer A (10 mM potassium phosphate buffer (pH 7.4) and 140 mM KCl). For guanidine titration, 5 μM protein was made with same buffer components but dissolved in 8 M guanidine hydrochloride. Protein-DNA complex midpoint melting temperatures (T_m_) were measured by CD at α-helical wavelength 222 nm. Protein-DNA complexes were made in Buffer A at 25 μM; data collected from 4-95 °C.

### Fluorescent Resonance Energy Transfer

A 15-base pair (bp) DNA duplex was fluorescently modified at the 5’-end of each strand to contain either: fluorescein (donor) or tetramethylrhodamine (TAMRA; acceptor). Kinetic *off*-rates (*k_off_*) were determined using stopped-flow FRET wherein a solution containing an equimolar labeled-DNA and protein was rapidly mixed with a solution containing an unmodified 15-bp DNA target. Donor emission was monitored at 520 nm, and *off*-rate constants determined using a single-exponential decay model. Specific DNA binding (K_D_) was determined using a constant concentration of modified DNA (25 nM) with increasing concentrations of the proteins. The change in the FRET signals with increasing protein concentration were plotted and fit as described ^56^.

### NMR Spectroscopy

Proteins were dissolved in a nitrogen-purged buffer containing 10 mM potassium phosphate buffer (10% D2O (pH 7.4)) and 50 mM KCl and placed in a 300-µL Shigemi NMR tube. Spectra of DNA and protein-DNA complexes were obtained at 700 MHz and 25 °C. Resonance assignments were obtained by analogy to previous NMR studies ^54; 57^.

### Protein modeling

Three-dimensional (3D) models of variant SRY HMG box/DNA complexes (containing specific recognition subsite 5’-TTTGTG-3’ and complement) were constructed based on the WT complex (Protein Databank [PDB] entry 1J46) and p.Met64Ile variant complex (PDB entry 1J47) as determined by Clore and colleagues using solution 3D/4D-NMR methods ^49^. Amino-acid substitutions were introduced and visualized using Pymol as described ^43; 57^. The protocol enabled adjustment of local dihedral angles to avoid steric clash; global molecular dynamics simulations were not performed.

## Results

### Genetic analysis

DNA sequencing of the *SRY* gene in the proband (P1) and her sister (P2), brother, father and grandaunt revealed a novel c.190A→G transition, predicting a p.Met64Val amino-acid substitution in the HMG box (Fig. 2). The same mutation was observed in the normal prepuberal male brother, fertile father, and a paternal 46, XY grandaunt (Fig. 1). Because of the chemical similarity of Met and Val as hydrophobic non-aromatic residues, complete genomic sequencing of P1, P2 and their father was undertaken to exclude DSD-associated mutations in other genes involved in gonadogenesis or sexual differentiation (Table 2). Two unannotated sequence variants (pThr239Pro and pLys242Gln) were identified in one *SOX9* allele in each individual. These variants have singly been observed in the general human population without a disease association, but to our knowledge their tandem occurrence has not previously been reported. Because mutations in *SOX9* can phenocopy Swyer mutations with respect to gonadal dysgenesis (typically in conjunction with abnormalities of cartilage and bone; campomelic dysplasia), functional studies of WT and variant SOX9 proteins were undertaken in cell culture (described below), providing evidence of neutral polymorphisms.

**Figure 2.**
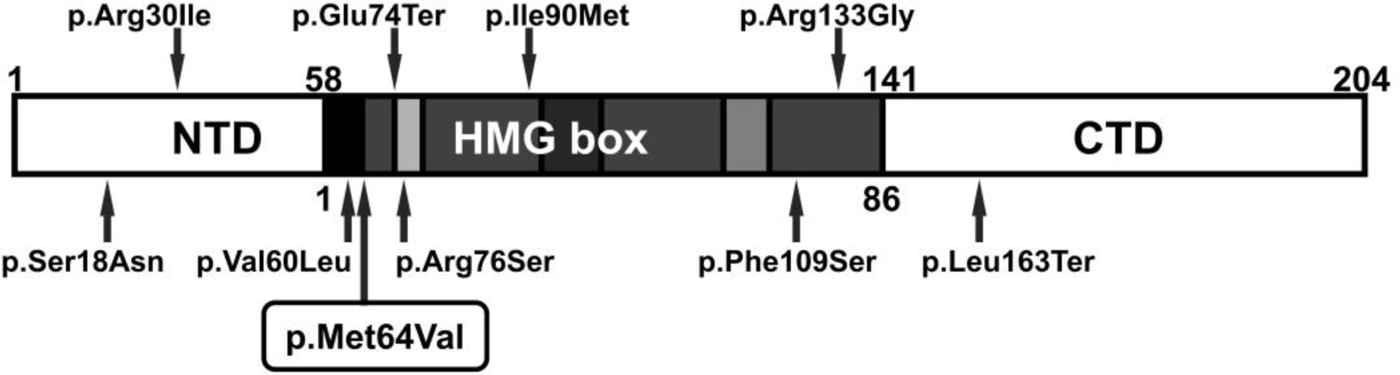
Broad distribution of non-mosaic inherited pathogenic variants reported in *SRY* gene. NTD: Amino terminal domain; CTD: Carboxyl terminal domain

### Gonadal histology assessment

Histopathologic examination of **P1** (Fig.3a) revealed ovarian stroma in the right gonad invaded by gonadoblastoma (GB); no follicles were observed. Her left gonad exhibited ovarian-type stroma and unidentified tubular features. Histopathologic findings of **P2** specimens were remarkable for bilateral neoplasia. The left Ov (Fig.3d-f) was grossly distorted with ovarian stroma invaded by both dysgerminoma (DG) and GB; the right Ov contained GB and highly fibrous ovarian structure with no demonstrable follicles. Some germ cells of both GB and DG exhibited nuclear expression of POU transcription-factor OCT3/4 (Fig. 3, b, e). In both sisters AMH/MIS, CYP11A1 and CYP19 (aromatase cytochrome P450) were expressed in the GB cells (Fig. 3, g-l).

**Figure 3.**
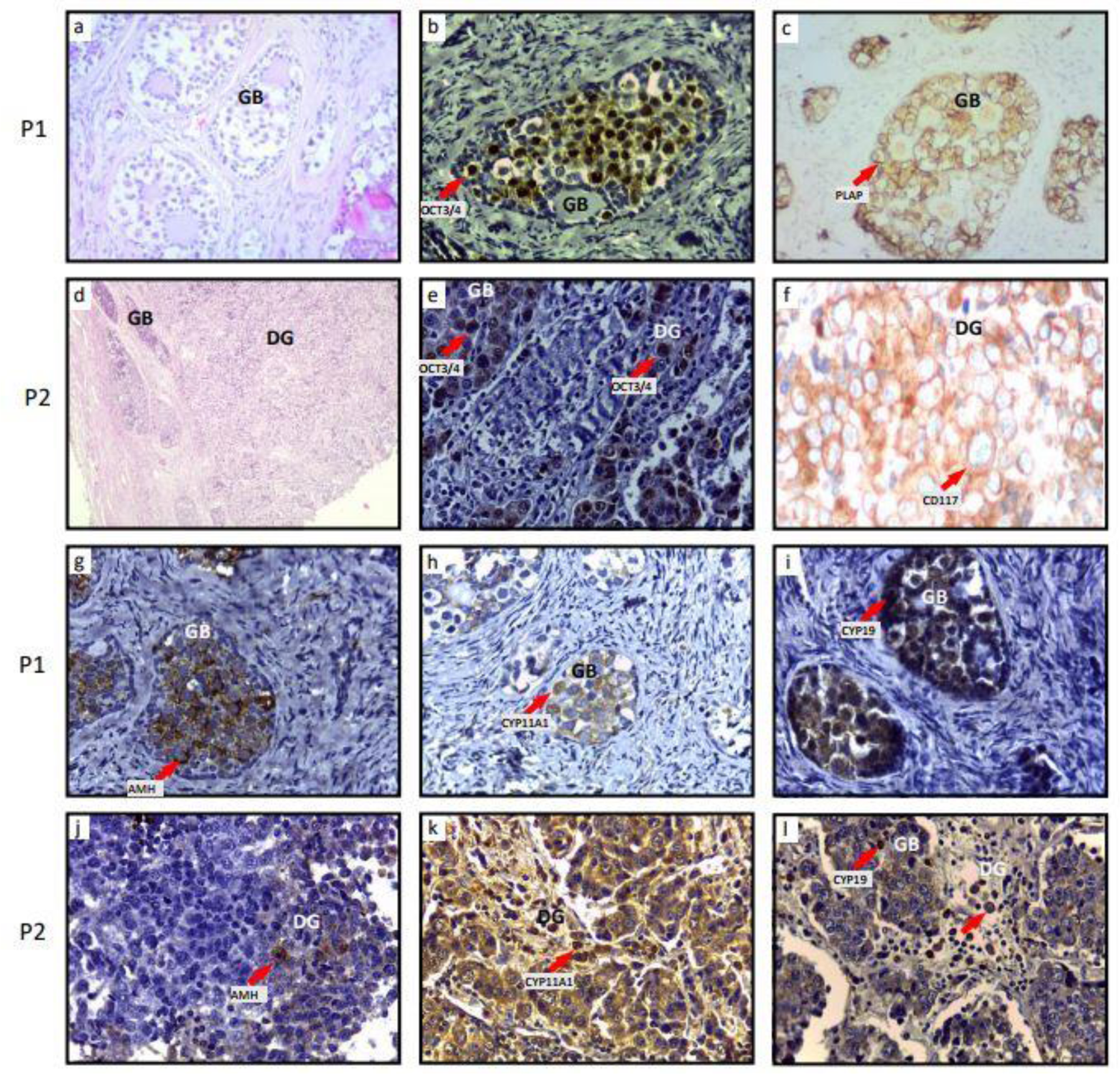
P1, right gonad (a, b, c) (a) Ovarian stroma with several nests of Gonadoblastoma (GB). No follicles were found, H&E, 20x; (b) OCT3/4 nuclear positive immunoexpression in germ cell of a GB, 40x; (c) PLAP positive cells in the nests of GB, 40x; P2, Left gonad (d, e, f) (d) Dysgerminoma (DG) and GB, H&E, 10 x; (e) OCT3/4 positive nuclear immunostaining in some cells of the DG and GB, 40x; (f) Cd117 positive cells in a DG, 40x. P1, right gonad (g, h, i); (g) AMH positive in cells of the GB, 40x; (h) CYP11A1 immunoexpression in cells of the GB, 40x; (i) CYP19 positive immunostaining in the GB, 40x; P2, Left gonad (j, k, l) (j) AMH positive immunostaining in some cells of the DG, 40x; (k) CYP11A1 positive cells inside the cell nests and outside them, 40x; (l) CYP19 positive immunostaining in small cells in GB and DG, 40x. Positive immuinostaining are depicted by red arrows.

### Functional Assessment of SOX9 Variants in Cell Culture

The two amino-acid substitutions found in one *SOX9* allele in P1, P2 and their father (pThr239Pro and pLys242Gln) were introduced, singly and together, into a SOX9 expression plasmid to enable functional assessment in a transcriptional reporter assay (Fig. 4a). Negative controls were provided by a SOX9 variant containing either (a) a deletion in its HMG box or (b) a point mutation in the HMG box (pMet113Ala), each known to block specific DNA binding ^58^. To monitor SOX9 activity, a luciferase reporter was constructed based on the well-characterized SOX9-responsive enhancer element of the murine *Col2a1* gene ^59; 60^; the transient co-transfection assays employed 293T cells. These studies demonstrated that transcriptional activity is not affected by substitutions pThr239Pro and pLys242Gln (individually or together) and so appear to represent neutral polymorphisms (Fig. 4b). These findings focused our attention on how SRY mutation pMet64Val might perturb its gene-regulatory function in accordance with the sex-reversed phenotypes of P1, P2 and other family members and yet also be compatible with the fertile male development.

**Figure 4.**
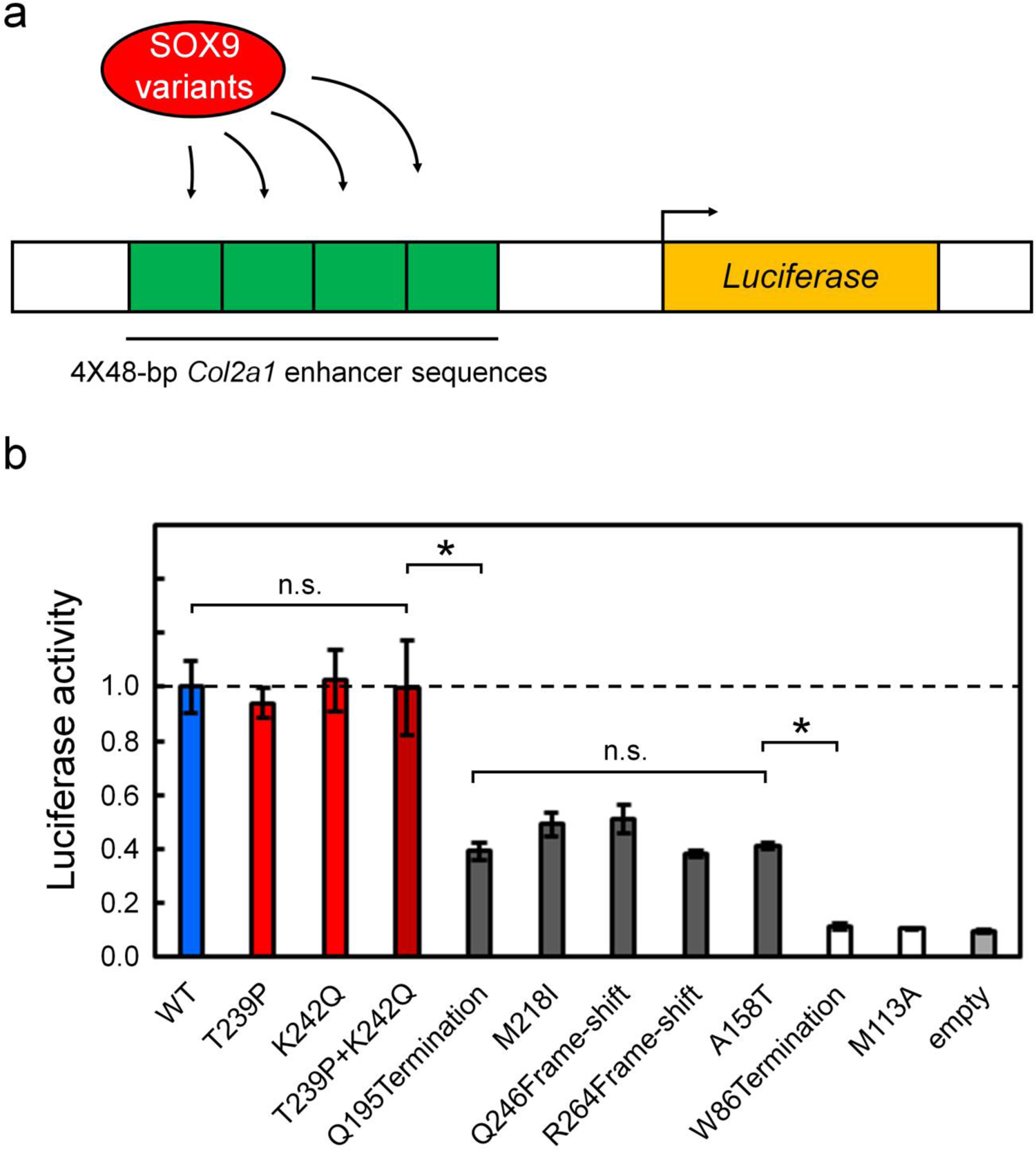
Luciferase-based co-transfection assay of SOX9 and variants in human 293T cell line. (a) Design of luciferase reporter construct. 4 of 48-bps elements derived from the enhancer region of *Col2a1* gene sites are shown as green boxes. Co-transfected SOX9 variants (red) would recognize this region and activate the luciferase reporter genes (yellow). Transfection of 293T cells with *SOX9* or variants (100 ng plasmid) stimulates expression of firefly luciferase. Assay was normalized by the co-transfected control Renilla plasmid. Mutations identified by the WGS result (T239P; red, K242Q; red, and double mutants; brown) were tested and exhibited WT-like luciferase activity (blue) whereas several clinical mutations show significant reduced activities (dark grey). Controls were provided by a truncated SOX9 mutant without HMG box (W86termination; white) and a mutation exhibiting abolished DNA-binding function (M113A; white). Empty coding-plasmid (light grey) shows the baseline readouts in this assay. Statistical comparisons: p-value (*) < 0.05; “ns” indicates p-value > 0.05.

### *In vitro* functional analysis of mutant p.Met64Val SRY

A rat pre-Sertoli cell line (designated CH34 cell) provided a model of the bipotential gonadal ridge ^4^, where in stage-specific SRY expression initiates male gonadogenesis via transcriptional activation of *SOX9* ^4; 42^. In this developmental program SRY binds within the core elements of two testis-specific enhancer elements (designated *TESCO* and *Enh13*, respectively; Fig. 5a) ^14; 16^. These regulatory events were recapitulated in the cell line, which was derived from the rat embryonic XY gonadal-ridge cell line just prior to its morphologic differentiation ^4; 10^.

**Figure 5.**
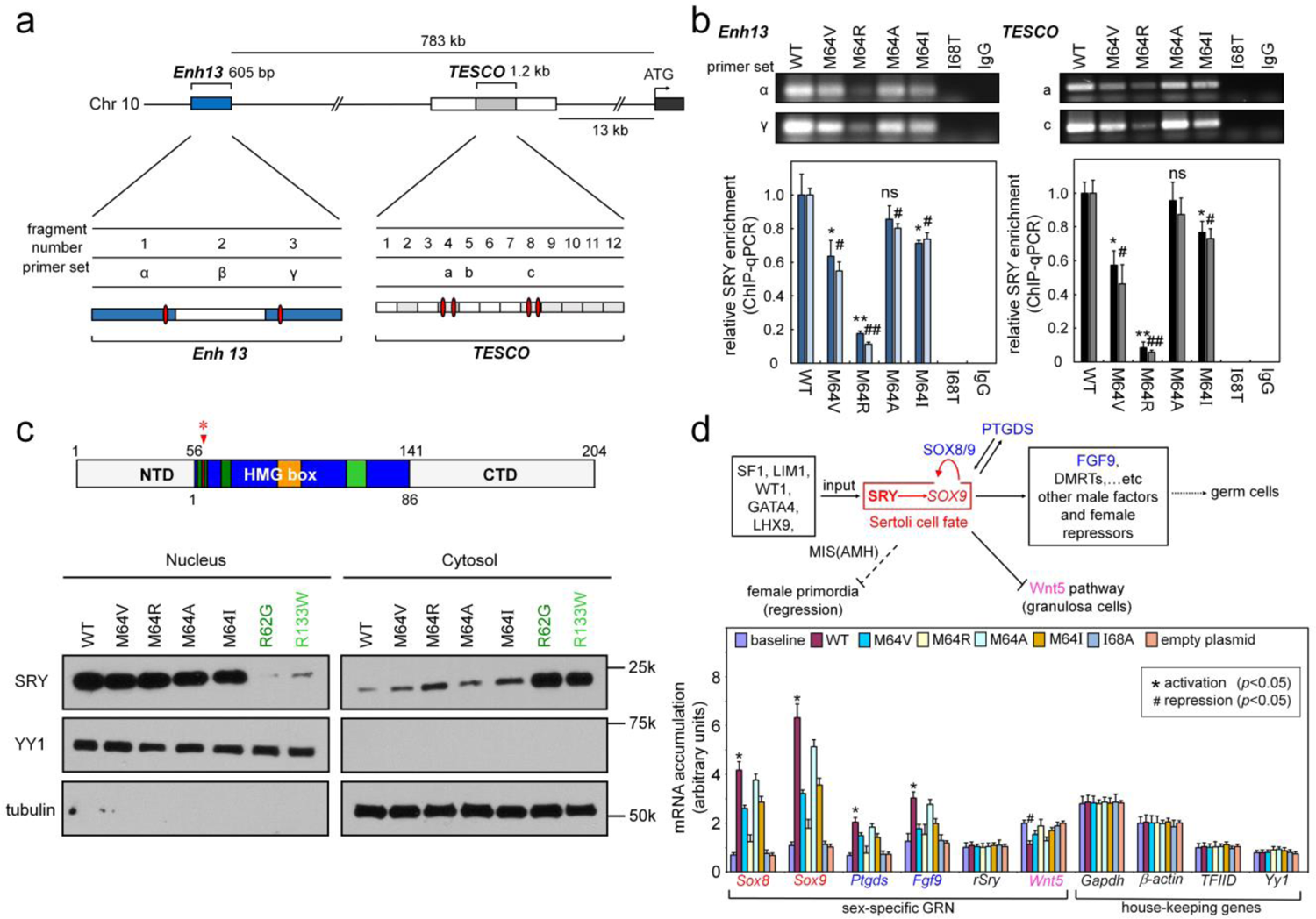
Analysis of SRY variants in a rat embryonic pre-Sertoli cell line. (a) *Sox9* gene is regulated by a far upstream enhancer element (blue: *Enh13* ^16^) and *TES* core elements (*TESCO*; light gray). Potential SRY binding sites are present in *Enh13* fragments 1 and 3 (related to primer sets α and β) and *TESCO* fragments 4 and 8 (primer sets a and c). (b) Effects of amino-acid substitutions on SRY enhancer occupancy. SRY-ENH13 (left) and SRY-TESCO (right) complexes were probed by anti-HA WB in ChIP assays (representative gel image in upper panel) and quantified by qPCR; results are summarized by histogram (lower panel). *Filled* and *open* bars represent occupancies of *Enh13* and *TESCO* fragments, respectively. Results were normalized relative to WT SRY (far left in each panel). Control lanes p.Ile68Thr and IgG indicate inactive clinical p.Ile68Thr SRY ^62^ and non-specific control. (c) *Top*, domain organization of human SRY. Residues 56-141 comprise the HMG box (blue) ^56^ (consensus numbers under the HMG box). Motifs within box: bipartite N-terminal nuclear localization signal (N-NLS; dark green) ^10; 86^, nuclear export signal (NES; orange) ^10^, and C-terminal NLS (C-NLS; light green) ^7^. Position Met64 is highlighted (red asterisk and arrow). *Bottom*, subcellular distribution of SRY variants was analyzed by immunoblotting in cytoplasmic or nuclear fractions. Controls were provided by NLS variants R62G and R133W (dark and light green, respectively) ^7; 65^. (d) Transcriptional survey of selected mRNA abundances following transient transfection of WT SRY or variants in gonadal lines CH34. Upper: Central SRY→*SOX9* regulatory axis (red box) with inputs in bipotential gonadal ridge (left box) leading to activation of male-specific program (right box) with repression of female development. Red curved arrow indicates SOX8/9 feedback, proposed to maintain *SOX9* expression following down-regulation of *SRY*. FGF9, fibroblast growth factor 9; GATA4, GATA binding protein 4; LHX9, LIM homeobox 9; LIM1, homeobox protein Lhx 1; MIS (AMH), Müllerian Inhibiting Substance (Anti-Müllerian Hormone); PTGDS, Prostaglandin D2 synthase; SF1, steroidogenic factor 1; WT1, Wilms’ tumor 1; Wnt5, wingless-type. Bottom: The baseline abundances in non-transfected cell, WT SRY, Met64 SRY variants, inactive control p.Ile68Ala SRY, and empty plasmid. Surveyed mRNAs include sex determination-specific genes *Sox*8, *Sox9*, *Ptgds*, *Fgf9*, *rat Sry*, and *Wnt5* ^55^. Controls were provided by housekeeping genes. *Inset box*, statistical significance at the level p<0.05 (Wilcox test) for SRY-dependent transcriptional activation (*) or repression (#).

Following transient transfection of WT or variant epitope-tagged SRY constructs (see Methods), SRY-directed transcriptional activation of the endogenous *Sox9* gene led to SRY occupancy of *TESCO* and *Enh13* as demonstrated by ChIP (Fig. 5b). Four substitutions were evaluated at SRY position 64 (consensus box position 9 in the hydrophobic wedge ^33; 49; 61^: p.Met64Val, related clinical variants p.Met64Ile and p.Met64Arg, and canonical Ala substitution p.Met64Ala. Negative controls were provided by an empty plasmid and by expression of p.Ile68Thr SRY ^62^, in which a known clinical mutation at the “cantilever” position of the SRY HMG box ^63^ markedly impairs its specific DNA binding ^64^. Additional controls were provided by well-characterized clinical variants within the nuclear localization signals of the HMG box (p.Arg62Gly ^65^ and p.Arg133Trp ^7^; Fig. 5c, lower panel); their impaired nuclear entry has previously been shown to attenuate *Sox9* enhancer occupancy and in turn decreased transcriptional activation ^10; 23; 35^. Similar results were obtained in a human prostate-cancer cell line (LnCaP; derived from a pulmonary metastatic deposit), suggesting that functional perturbations caused by Met64 mutations in a rat pre-Sertoli-derived cell line are independent of species.

In CH34 cells the SRY variants at position 64 exhibited, to varying extents, reduced *TESCO*/*Enh13* enhancer occupancies. Whereas canonical p.Met64Ala SRY preserved >80% of occupancy relative to WT (as quantified by ChIP-qPCR), for example, *de novo* clinical variant p.Met64Arg exhibited only 20% occupancy (Fig. 5b). Attenuation due to p.Met64Val orp.Met64Ile was intermediate (*ca*. 60 and 70%, respectively). As expected, p.Ile68Thr blocked detectable binding of the variant SRY at the *TESCO* and *Enh13* enhancers. Subcellular fractionation studies (validated by respective nuclear and cytoplasmic markers YY1 and tubulin ^10^) demonstrated that—with the exception of the NLS control variants—the mutant SRYs exhibited predominantly nuclear localization in accordance with the WT pattern (Fig. 5c). The gene-regulatory activities of Met64 SRY variants, evaluated via the pivotal SRY/*Sox9* axis (red box in upper panel of Fig. 5d), were likewise attenuated to varying extents as evaluated by qPCR (at left in Fig. 5d, lower panel). Strikingly, among the SRY variants, the degree of impaired transcriptional activation correlated with the extent to which enhancer occupancies were attenuated. Reduced transcriptional activation of *Sox9* was associated with concordant defects in the activation of autosomal genes downstream of *Sox9* in the male program (*Sox8*, *Ptgds* and *Fgf9*); the expression of house-keeping genes (*Gapdh*, β*-actin*, *tbp* and *yy1*) was by contrast not affected (at right in Fig. 5d, lower panel).

### Biophysical analysis and molecular modeling

How pMet64Val affects the DNA-binding and DNA-bending properties of the SRY HMG box was investigated by fluorescent methods ^54; 57^. These studies employed a 15-base pair (bp) DNA duplex in which the 5’-end of one strand was flexibly linked to a fluorescence donor (fluorescein) whereas the 5’-end of the other strand was flexibly linked to a fluorescence acceptor (tetramethylrhodamine; TAMRA). Binding of the SRY HMG box to the central DNA target site (5’-ATTGTT and complement) leads to bending-induced reduction in the end-to-end DNA distance (Fig. 6a). This change in distance may in turn be monitored by FRET enhancement (Fig. 6b). Such spectroscopic studies demonstrated that binding of the WT and variant HMG box was associated with indistinguishable FRET enhancements (black and red spectra in Fig. 6b) relative to the free (unbent) DNA duplex (green spectrum). This system enabled determination of the specific protein-DNA dissociation constant based on quantitative analysis of FRET efficiency as a function of protein concentration (Fig. 6c). The affinity of the variant domain (K_D_ 32 nM) is slightly weaker than that of the WT domain (K_D_ 24 nM) as summarized in Table 3. Despite this modest change, stopped-flow kinetic studies uncovered a more substantial perturbation: the *off*-rate (*k*_off_) of the pMet64Val domain, reflecting the kinetic lifetime of the variant protein-DNA complex, was accelerated by more than threefold relative to the WT complex (Fig. 6d and Table 3). Such kinetic instability was associated by subtle changes in the ^1^H-NMR spectrum of the variant domain-DNA complex (Fig. 7), as monitored via the imino Watson-Crick resonances of thymidine and guanine. Whereas titration of the WT domain is in slow exchange on the time scale of NMR chemical shifts, the variant spectra continues to exhibit discrete free and bound imino resonances but with subtle broadening indicating an increase in exchange rate between free and bound DNA states. Conformational broadening is especially prominent in the indole NH resonances of major-wing core residues Trp70 and Trp98 (asterisks in Fig. 7; consensus box positions 15 and 43, respectively), indicating that the bound state of the variant domain is molten or in intermediate exchange between conformational substates. The indole resonance of surface residue Trp107 (box position 52) by contrast remains sharp, reflecting motional narrowing as on the surface of the WT domain.

**Figure 6.**
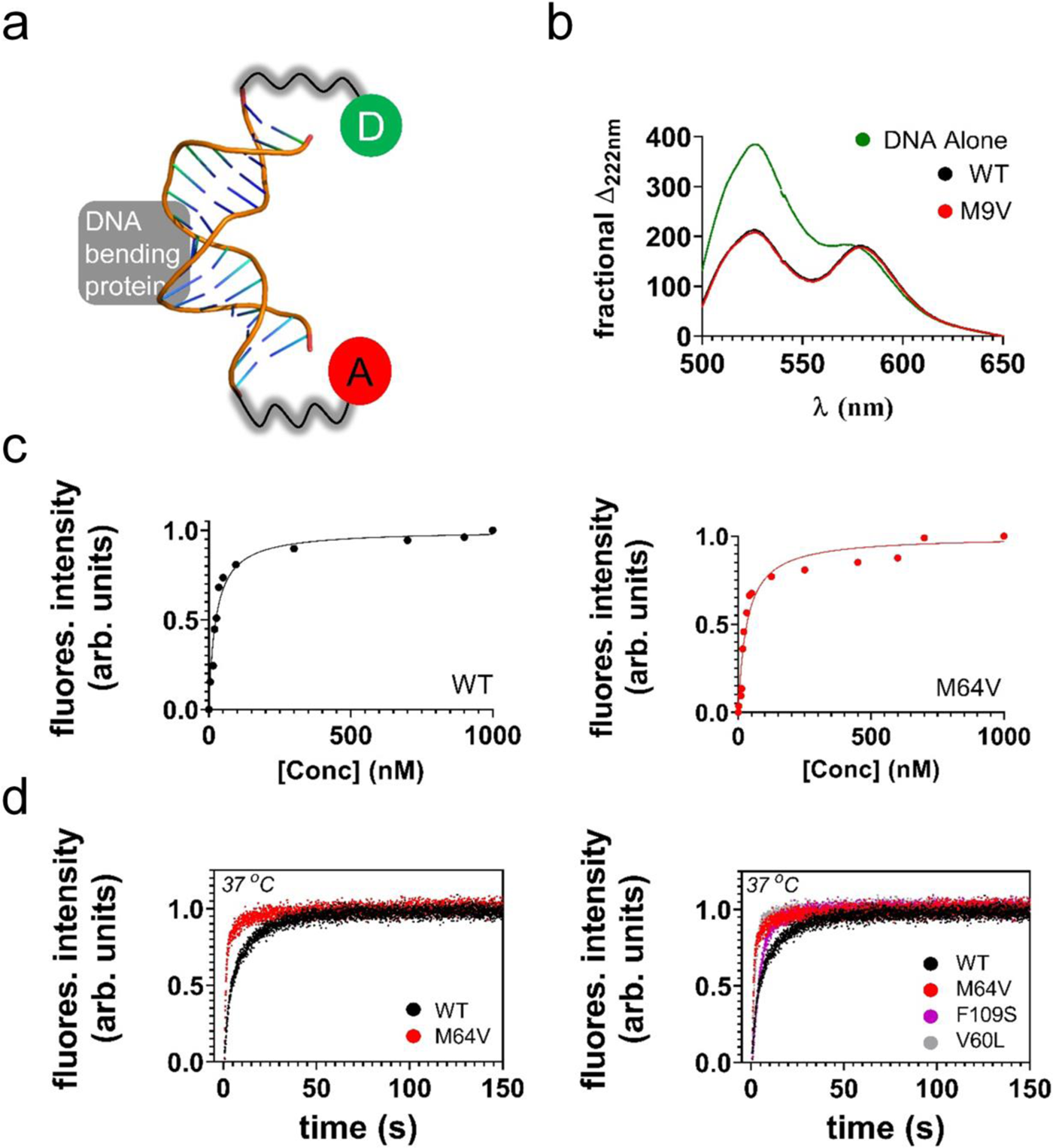
(a) Representative picture of the DNA used in FRET-based assays. A 15bp duplex is modified at the 5’-ends of each strand with a donor (D, fluorescein, green) and acceptor (A, TAMRA, red). DNA model has been modified from published SRY-DNA complex ^49^. (b) steady state FRET measurements indicating that the bent protein-DNA architecture between the WT and M9V mutant domain is similar. (c) Binding affinity of the WT and pMet64Val mutant HMG domains determined by FRET. The mutant domain (red trace) exhibits a small decrease in DNA binding at physiological temperature (37 °C). (d) Stopped flow FRET assay in the determination of the off rate for the WT and mutant domain at physiological temperature. The mutant domain (red trace) has an increased off rate compared to WT (black trace). The off rate of the mutant domain is similar to kinetic defects of other inherited clinical SRY mutations (F109S purple trace and V60L grey trace)

**Figure 7.**
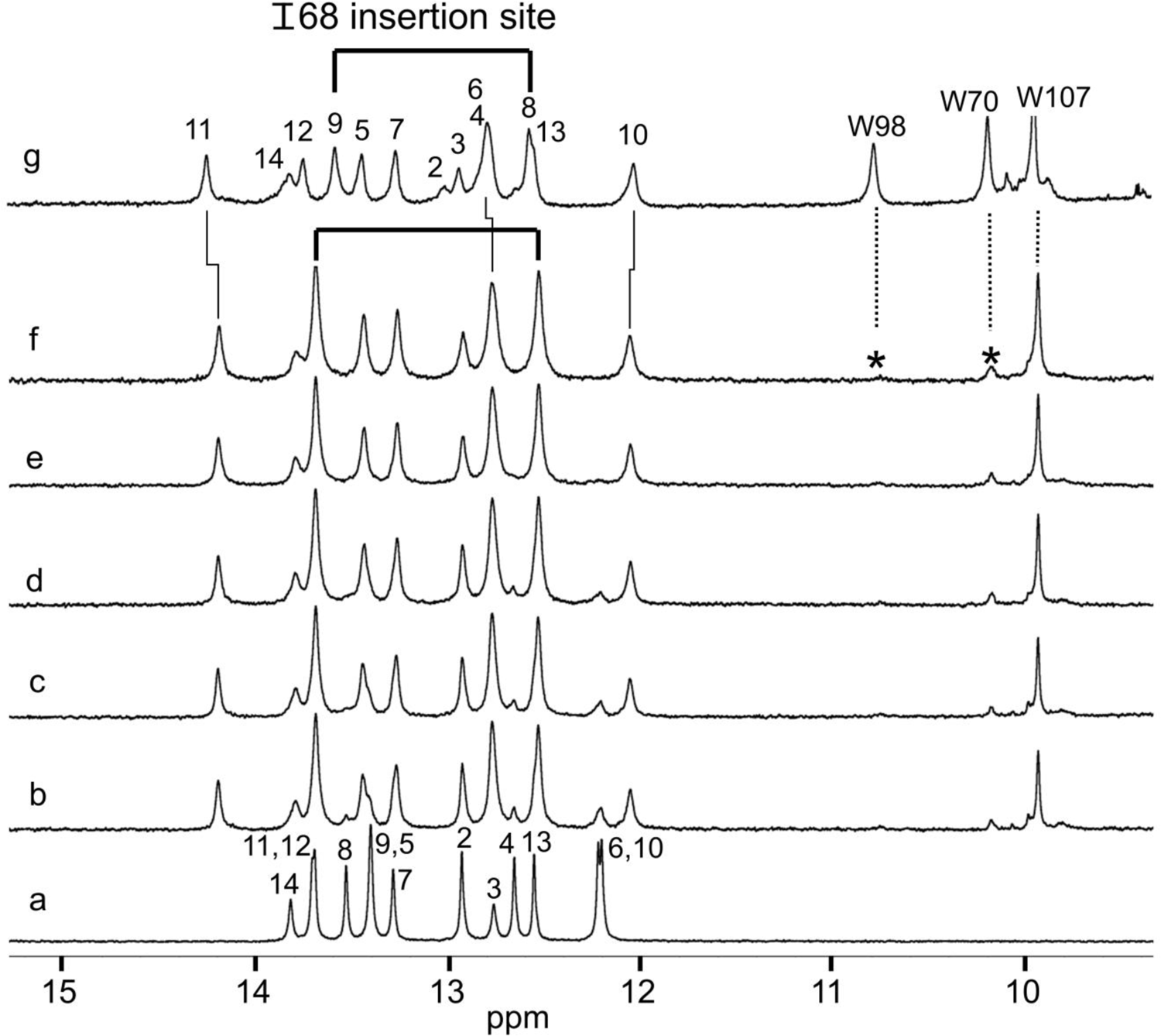
1D ^1^H-NMR protein-DNA titration. Assignments are as indicated (numbering scheme; top): spectra of free DNA (a), native equimolar SRY-DNA complex (g) and variant-equimolar DNA complex (f); respective spectra b–e were obtained on addition of successive aliquots of the variant SRY domain to obtain protein-DNA stoichiometries are 1:0.2, 1:0.4, 1:0.6 and 1:0.8. Vertical segments between spectra f and g indicate small differences in chemical shifts; the trend is toward attenuated complexation shifts in the variant complex. Horizontal bracket at top indicates side-chain insertion between bp 8 and 9 (partial intercalation) by “cantilever” residue Ile-68; see Figure 8 for diagnostic intermolecular NOEs. Asterisk in panel f highlights line broadening of the indole NH resonances of Trp70 and Trp 98; the corresponding indole resonance of Trp107 (on the protein surface) exhibits motional narrowing in both the WT and variant spectra. Imino resonances of base pairs 1 and 15 are not seen due to fraying; the resonances of base pairs 2 and 14 are broadened.

**Table 3.** DNA binding constants and *off*-rates

| Protein | $K_D$ (nM)<sup>a</sup> | $k_{off}$ (s<sup>-1</sup>) | lifetime (ms)<sup>b</sup> |
| --- | --- | --- | --- |
| WT | 24.7 ±2.3 | 0.122 ±0.009 | 8.2 x 10<sup>3</sup> |
| M64V | 32.3 ±4.0 | 0.409 ±0.01 | 2.4 x 10<sup>3</sup> |
| V60L | 32.2 ±2<sup>1</sup> | 0.426 ±0.01 | 2.3 x 10<sup>3</sup> |
| F109S | 26 ±6<sup>2</sup> | 0.193 ±0.008 | 5.2 x 10<sup>3</sup> |

<sup>a</sup> Values were determined using HMG-box fragments at physiological temperature (37 °C)
<sup>b</sup> determined as $1/k_{off}$
<sup>1</sup> previously reported <sup>54</sup>
<sup>2</sup> previously reported <sup>43</sup>

The mutation does not prevent partial intercalation of the “cantilever” side chain Ile68 (box position 13) between AT base pairs as indicated by the signature chemical shifts of the flanking thymidine imino resonances (brackets in Fig. 7; ^62^) and by intermolecular nuclear Overhauser effects (NOEs) from these DNA protons to the δ-methyl group of the Ile68 side chain (Fig. 8; ^62^). Nonetheless, the detailed pattern of DNA complexation shifts (vertical lines in Fig. 7 and Table 4) reveals a trend toward attenuation of the magnitude of such sifts relative to the free DNA site. Such subtle attenuation in the spectrum of the variant complex suggests greater conformational averaging of bound-state chemical shifts. Such averaging can be due to enhanced local structural fluctuations ^56^.

**Figure 8.**
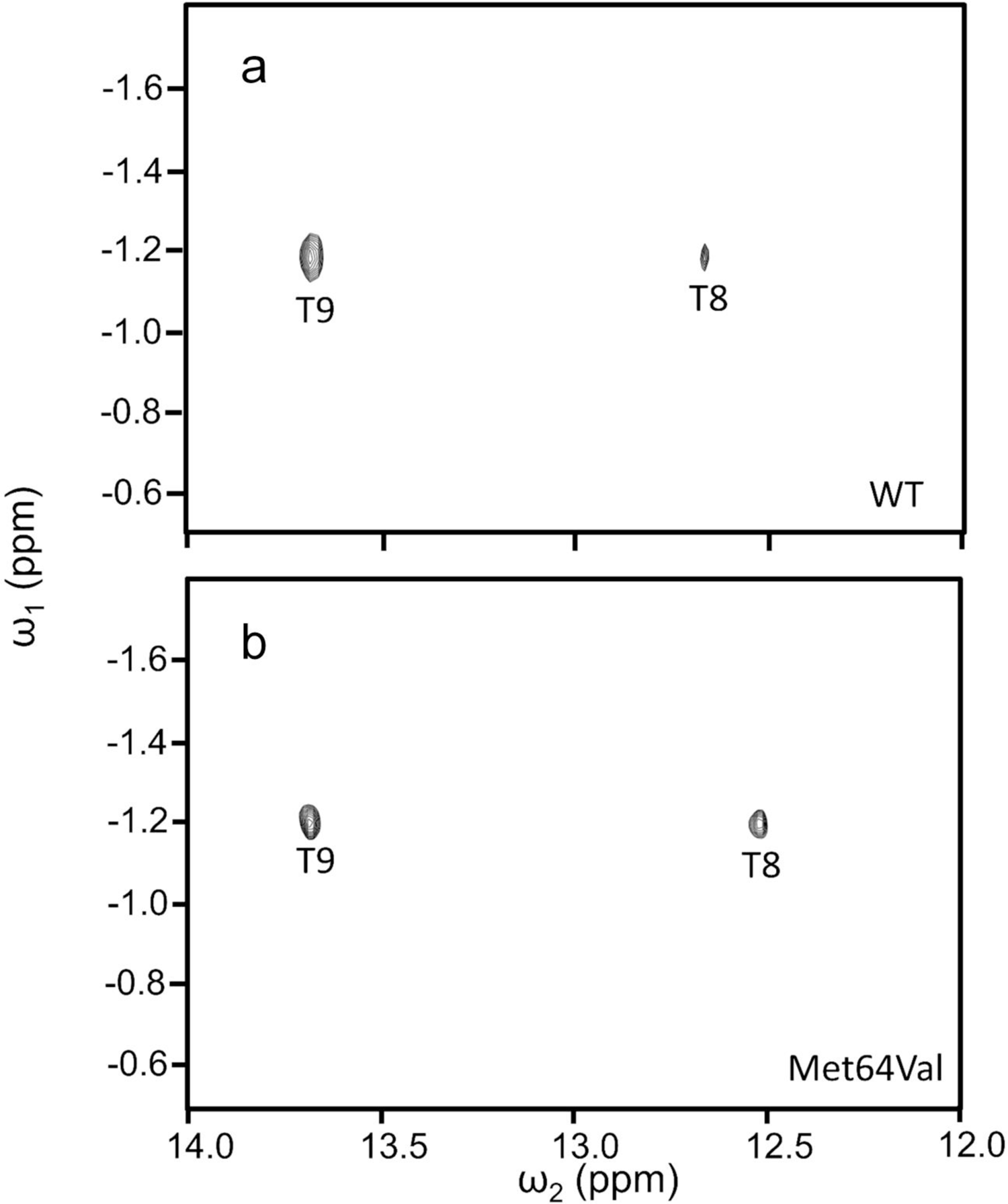
2D ^1^H-NMR NOESY cross peaks diagnostic of SRY-DNA intercalation. Spectra of (a) WT SRY HMG box/DNA complex and (b) Met64Val SRY HMG box/DNA complex (conditions as in Fig. 7). Corresponding intermolecular NOEs are observed between the δ-methyl resonance of Ile68 (the cantilever side chain; vertical axis) and the flanking thymidine imino protons (horizontal axis) at the site of partial intercalation (bold in 5’-A**TT**GTT-3’ and complement; base pairs 8 and 9 in the 15-bp DNA duplex). The mixing time was in each case 150 ms. Spectra were obtained at 25 °C in 10 mM potassium phosphate (pH 7.4) and 50 mM KCl in 90% H_2_O and 10% D_2_O.

**Table 4.** DNA Imino ^1^H-NMR Chemical Shifts^a^

| base pair | free | WT | $\Delta\delta_{\text{hum}}$ | variant | $\Delta\delta_{\text{variant}}$ | $\Delta\Delta\delta$ |
| --- | --- | --- | --- | --- | --- | --- |
| 2<sup>b</sup> | 12.93<sup>c</sup> | 13.11 | 0.18<sup>d</sup> | 12.92 | -0.01<sup>d</sup> | 0.19 |
| 3 | 12.76 | 13.04 | 0.28 | 12.92 | 0.16 | 0.12 |
| 4 | 12.66 | 12.89 | 0.23 | 12.77 | 0.11 | 0.12 |
| 5 | 13.4 | 13.54 | 0.14 | 13.44 | 0.04 | 0.10 |
| 6 | 12.22 | 12.89 | 0.67 | 12.77 | 0.55 | 0.12 |
| 7 | 13.29 | 13.36 | 0.07 | 13.27 | -0.02 | 0.09 |
| 8 | 13.53 | 12.67 | -0.86 | 12.53 | -1.00 | 0.14 |
| 9 | 13.40 | 13.68 | 0.28 | 13.68 | 0.28 | 0.00 |
| 10 | 12.20 | 12.13 | -0.07 | 12.05 | -0.15 | 0.08 |
| 11 | 13.71 | 14.35 | 0.64 | 14.19 | 0.48 | 0.16 |
| 12 | 13.70 | 13.85 | 0.15 | 13.68 | -0.02 | 0.17 |
| 13 | 12.55 | 12.65 | 0.10 | 12.53 | -0.02 | 0.12 |
| 14 | 13.82 | 13.93 | 0.11 | 13.79 | -0.03 | 0.14 |

<sup>a</sup>NMR studies employed DNA duplex 5'–GGGGTG**ATTGTT**CAG-3' and complement; the core target site is in bold.
<sup>b</sup>Numbering scheme refers to top strand; primes (e.g., 7') refers to nucleoside of partner strand.
<sup>c</sup>Chemical shifts are given in parts per million (ppm) relative to DSS, assumed to be at 0 ppm.
<sup>d</sup>Positive values of $\Delta\delta$ indicate upfield shifts in complex (i.e., to lower ppm values); negative values represent downfield shifts.

The conservation and structural environment of the variant p.Met64Val side chain was visualized in a rigid-body model derived from the WT solution structure of the SRY HMG box/DNA complex (Fig. 9) ^49^. Residue 64 (box position 9; Fig. 9a) exhibits similar side chain orientation in deposited structures as well as comparable *ca.*x^1^ and x^2^ angles (averaged x^1^ = −75.4° ±-9.0 x^2^ = −171.3 ° ±-4.9, Fig. 9b). Met64 is part of a “hydrophobic wedge” of side chains (Fig. 9c) that engages the minor groove of the bent DNA site (positions 67, 68 and 98; purple in Fig. 9c). Such non-polar engagement is central to the mechanism of sharp DNA bending. The aliphatic portion of the Met64 side chain and its large sulfur atom project a top the “cantilever” side chain I68 (box position 13) to pack near the deoxyribose moiety of an adenosine (bold in CACAA**A**A-5’) (Fig. 9d). A Cori-Pauling-Kolton (CPK) hard-sphere model of the interface highlights the space-filling role of the Met64 side chain at one edge of the wedge deformation (Fig. 9d). In addition, the sulfur atom makes a hydrogen bond with the guanidine group of Arg72 (Fig. 9c, d; in gold) ^48^. The side chain of Arg72 makes contacts with the DNA but is partially exposed in the bound structure. Substitution of Met64 by Val appears readily accommodated. Although Val is smaller than Met and different in shape, no steric clash would be introduced in an appropriate x_1_rotamer. In the context of the WT SRY-DNA complex, the substitution might nonetheless be associated with a local crevice (Fig. 9e, f). It is possible that the distinct size and shape of Val may require subtle adjustments in the neighboring DNA conformation as illustrated by the solution structure of a p.Met64Ile variant complex (albeit with variant DNA core target site 5’-TTTGTG-3’ and complement; see Discussion) ^49^.

**Figure 9.**
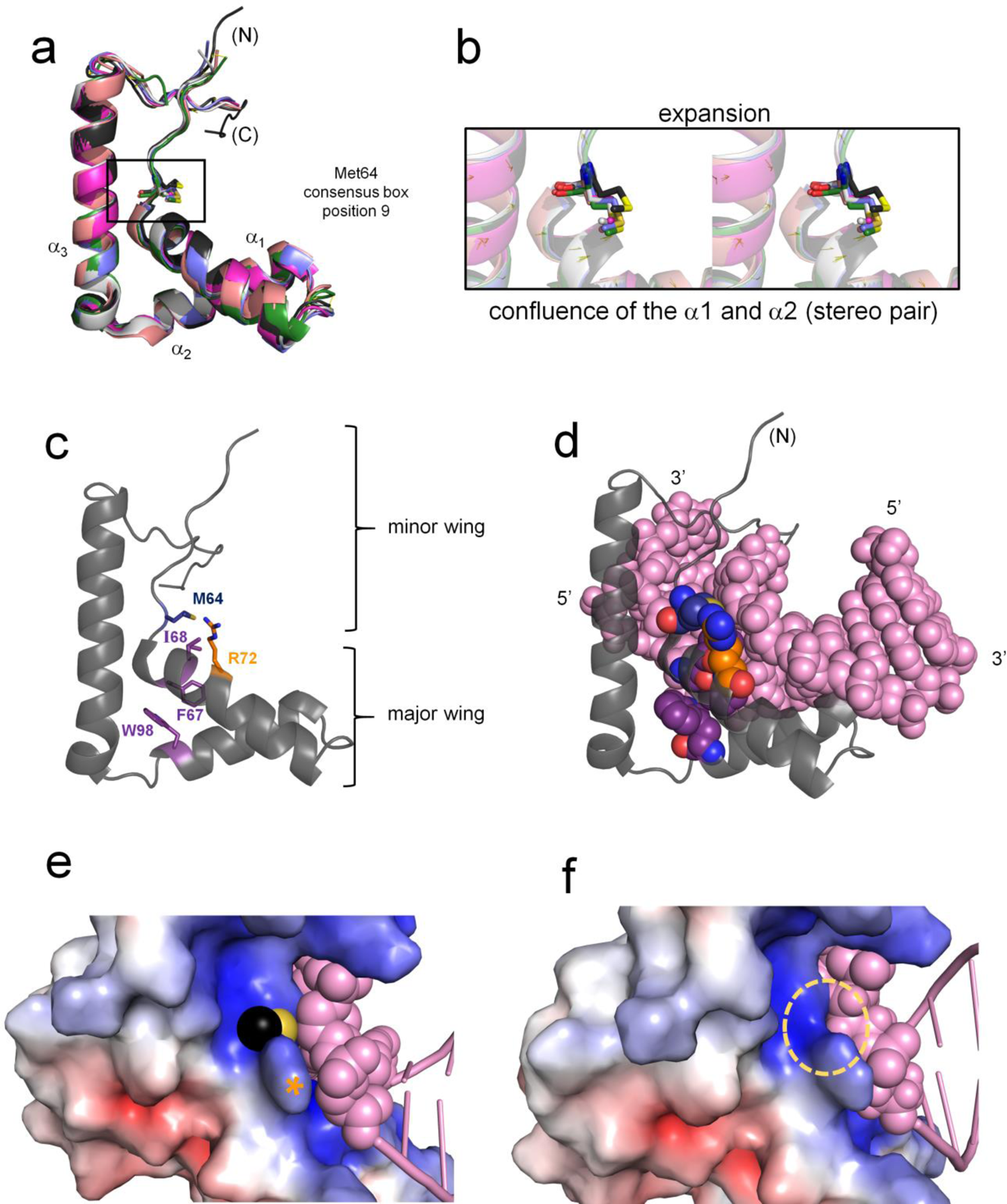
Structural modeling of p.Met64Val SRY HMG box. (a) Alignment of SRY and SOX HMG-box domains (as deposited in the Protein Data Bank [PDB]; see below). The ribbon models highlight structural conservation of an L-shaped domain with three helices (α_1_, α_2_ and α_3_); also shown are the side chains of Met64 in human SRY (black) and homologous residues in SOX domains (consensus box position 9 as defined by Clore and colleagues ^87^). The structures were in each case extracted from specific DNA complexes. (b) Expansion of the aligned structures (box in panel a) highlighting similar orientation of the Met side chain and positioning of the sulfur atom (yellow) among structures shown in stereo. (c) Human SRY HMG box is shown as a grey ribbon. Met64 is part of a hydrophobic wedge motif that underlies the bent DNA surface (DNA not shown) side chains of the residues of this motif are labeled and shown in purple. Arg72 (orange) is the “nearest neighbor” to the Met side chain in structure. (d) SRY HMG box (gray ribbon) bound to a specific DNA site (pink CPK representation with 5’and 3’-ends as labeled). “Hydrophobic wedge” side chains are shown as CPK models in blue (Met64) or purple (Phe67, Ile68 and Trp98; box positions 12, 13 and 43) at or near the surface of an expanded DNA minor groove; Arg72is shown in gold. (e) Electrostatic surface of the WT SRY HMG box showing terminal atoms of Met64; the C_ε_ of Met64 is shown in black whereas its neighboring sulfur atom is yellow. (f) The putative crevice that would form on *in silico* deletion of the Met64 side chain in a rigid-body model (dashed yellow circle). Due to the small size of Val relative to Met, the variant is predicted to exhibit a packing defect at this surface. Color code in panel a: black, human SRY (PDB entry 1J46); pink, mouse Sox 4 (3U2B); green, human SOX9 (4EUW); light grey, mouse Sox18 (4Y60); lavender, mouse Sox9 (4S2Q); magenta, mouse Sox17 (3F27); and yellow, human SOX2 (1GT0).

## Discussion

Herein we have described clinical-pathological features associated with a novel inherited mutation in SRY encoding a substitution in its HMG box (p.Met64Val).This position is invariant in both Sry and an extensive metazoan SOX family of SRY-related HMG boxes ^62^. Cell-based studies of p.Met64Val SRY were undertaken in relation to WT SRY and *de novo* Swyer mutations at this position (p.Met64Ile ^50^ and p.Met64Arg ^51; 66^). In an embryonic pre-Sertoli XY cell line ^10^ and in a human cancer cell line the clinical mutations impaired SRY-mediated transcriptional activation of target gene *Sox9* in association with decreased occupancy of testis-specific enhancer elements *EHN13* and *TESCO*. Whereas p.Met64Arg resulted in marked gene-regulatory perturbations, p.Met64Val and p.Met64Ile were compatible with partial maintenance of such activity. Our *in vitro* results thus rationalize both the pathogenicity of p.Met64Val SRY in the probands and its continued biological function in their father and brother. The range of phenotypes in this family is reminiscent of the murine intersexual phenotypes associated with Y:autosome incompatibility ^67^.

### Family history and clinical features

The present pedigree contains two 46, XY sisters (probands P1 and P2), their XY father, prepuberal XY brother and an affected paternal XY grandaunt. Observation of the same *SRY* variant in an affected grandaunt makes paternal mosaicism unlikely. As in other Swyer cases ^68^, primary amenorrhea was observed in association with hypergona (dotropic hypogonadism due to estrogen deficiency. Unlike prior reports, the sisters exhibited breast development. Whereas the complete androgen insensitivity syndrome (due to androgen receptor mutations; “testicular feminization”) is a more frequent cause of 46, XY sex-reversal in conjunction with primary amenorrhea yet retained breast development ^69^,this etiology was excluded by both the absence of high serum testosterone and absence of maternal inheritance.

Gonadal tumors were found in the probands (gonadoblastoma in P1 and dysgerminoma/gonadoblastomain P2). In accordance with other reports ^70–72^, the presence of positive steroidogenic markers (such as CYP11A1 and CYP19) may rationalize the observed breast development and pubertal uterus size. Tumor steroid production might also have increased skeletal bone maturation, accounting for the heights of the sisters (shorter than expected in patients with 46, XY CGD ^68^).The presence of pluripotential aberrant germ cells was revealed by high nuclear OCT3/4 immuno-expression. These findings are in accordance with previous reports that 46, XY dysgenetic gonads have a high risk of developing germ-cell tumors ^73; 74^. Thus, aspects of the sisters’ phenotype may represent paradoxical normalization of pubertal development as a paraneoplastic syndrome. Similarly, spuriously normal AMH/MIS levels were presumably due to secretion by immature neoplastic cells (i.e., unrelated to its meaning as a marker of ovarian reserve in reproductive-age 46, XX females) ^75^.

Mitigation of the classical Swyer phenotype (increased stature with minimal breast development) may in principle be associated with (a) genetic mosaicism leading to distinct regions of ovarian or testicular differentiation, (b) rests of ovarian follicles in a non-mosaic ovotestis or (c) hormonal secretion by tumor stroma in a dysgenetic gonad. The first mechanism was exemplified by pLeu101His in mosaic 46, XY patient with atypical genitalia and ovotesticular 46, XY DSD ^76^. This mutation presumably destabilizes the core of the SRY HMG box. The second mechanism was reported in a 46, XY infant with female external genitalia but no Müllerian structures ^21^; the gonads were partially descended with ovotestis on one side. The mutation (p.Val60Ala) causes a subtle destabilization of the box’ minor wing ^54^ and partial impairment of nuclear import ^10^. Because the gonads were removed in infancy as part of surgical assignment of female sex, pubescent development of secondary sexual characteristics could not be evaluated. Secretion of sex steroids by the stromal component of gonadoblastoma in association with breast development was first suggested by Vilain and colleagues in 2002 in a 46, XY patient with Swyer syndrome ^38^. The mutation (p.Tyr127Phe) represents a subtle modification of the protein-DNA interface. Although reported to abolish specific DNA binding, in our hands substantial activity is retained *in vitro* (unpublished results).

### Functional studies in an embryonic XY pre-Sertoli cell model

The functional consequences of p.Met64Val and other mutations at this site in the SRY HMG box were investigated in a rodent cell line (CH34) derived from the embryonic XY gonadal ridge just prior to its morphologic differentiation ^4^. Presumed to model the human pre-Sertoli cell at the site and stage of SRY’s regulatory function in testis determination ^4^, CH34 cells provide an epigenetic milieu in which WT SRY can function as a gene-specific transcriptional activator of endogenous target gene (*Sox9*) even when expression of the transfected protein is attenuated to 10^2^-10^4^ protein molecules per cell (by dilution of the SRY plasmid by a fifty-fold excess of the empty plasmid ^10^). As described in previous studies ^10^, key validation of this model was provided by several criteria: (i) enhanced Sox9 mRNA accumulation on transient transfection of WT SRY but not on transient transfection of diverse clinical SRY variants associated with *de novo* somatic sex reversal ^10; 43; 57^; (ii) ChIP-based observation of SRY localization to testis-specific enhancer elements upstream of the Sox9 transcriptional start sites and not at other potential enhancer elements in this highly regulated gene; (iii) selective transcriptional activation of a downstream pathway including fetal testis-specific Sox9 targets (*Sox8*, *Amh/Mis*, *Ptgds* and *Fgf9*) ^43; 55; 77^; (iv) in the pre-Sertoli cell line, SRY-induced down regulation of the ovarian-specific pathway otherwise characteristic of granulosa cells ^10; 55^; and (v) absence of a nonspecific state of transcriptional activation or changes in expression of other SOX genes not pertinent to gonadal differentiation. Cell lines not fulfilling these criteria provided controls to distinguish the gene- and lineage-specific regulatory function of SRY from general effects potentially observed in cell culture on transient transfection of an architectural transcription factor. Control studies of human XY LnCaP cells (derived from metastatic prostate cancer) suggested that species of origin did not confound the CH34 studies. The appropriateness of studying human SRY variants in a rat cell line is broadly supported by the native activity of a human SRY transgene (under the regulatory control of mouse control sequences) to trigger testicular differentiation (and male downstream development) in a XX mouse ^78–80^).

The partial biological activity of p.Met64Val SRY (*ca*. 50% with respect to both *Sox9* enhancer occupancy and *Sox9* transcriptional activation; Fig. 5b, d) rationalizes the phenotype of the father. That such a subtle decrease in activity can also lead to gonadal dysgenesis is consistent with previous studies of inherited Swyer alleles ^10; 43^. The tenuous threshold of male sex determination has also been observed in mouse genetics ^81^. Swyer allele p.Met64Ile exhibited an even more subtle perturbation (*ca.* 70% with respect to *Sox9* enhancer occupancy and 60% with respect to *Sox9* transcriptional activation; it is difficult to interpret this small but reproducible discordance between percent impairment of enhancer occupancy versus transcriptional activation as the former studies technically required SRY over-expression with 10^6^ molecules per cell). Although the p.Met64Ile mutation was uncovered as a *de novo* mutation in one small family ^50^, this observation does not in principle exclude its potential inheritability in other, more extensive pedigrees. The precise molecular threshold in human gonadogenesis governing the specification of maleness is likely to depend on genetic background as in mouse strains ^67^. *De novo* mutation p.Met64Arg led to a *ca*. tenfold reduction in enhancer occupancy (despite SRY over-expression) with complete abolition of *Sox9* transcription activation. Such a marked perturbation is in accordance past biochemical studies of the bending-defective p.Met64Arg HMG box ^52^.

Why p.Met64Val and p.Met64Ile impair enhancer occupancy is not well understood. Although defective DNA bending was initially ascribed to p.Met64Ile SRY ^82^, this finding was due to truncation of the C-terminal tail of the HMG box ^56; 61^. The complete motif (as well as the intact protein) exhibited native bending of a consensus DNA site (5’ATTGTT-3’ and complement) ^61^. It is possible that small changes in DNA bend angle, as observed in complexes with non-consensus DNA sites by permutation-gel electrophoresis ^61^, could lead to decreased enhanceosome assembly. Such altered bending was observed by NMR in studies of a lower-affinity DNA site (5’-TTTGTG and complement) ^49^. Testing this hypothesis will require fine mapping of SRY binding sites in the testis-specific enhancer elements of *Sox9* (*Enh13* and *TESCO* ^14; 16^) and their functional validation ^83^. Our present results based on biochemical subcellular fractionation do not support a previous hypothesis that p.Met64Ile impairs nuclear localization ^61^.

Although pMet64Val leads to kinetic instability of the variant HMG box/DNA complex (Fig. 6c and Table 3), it is not clear that this perturbation is sufficiently marked to affect transcriptional potency. Indeed, previous studies of unrelated inherited Swyer mutations pVal60Leu (V60L) and pPhe109Ser (F109S) also uncovered accelerated *off*-rates with modest reductions in affinity (Table 3), and yet under appropriate experimental conditions these variants are fully active in CH34 cells. (Assessment of pVal60Leu SRY required rescue of its nuclear localization by fusion of a nuclear localization signal [NLS; from SV40]^10^ whereas assessment of pPhe109Ser SRY required chemical proteosome inhibitor to mitigate accelerated intracellular degradation ^43^.) It would be of future interest to investigate sites of chromosomal binding of the WT and variant SRY proteins via ChIP-Seq and ChIP-exo methods to obtain a bias-free and genome-wide perspective.

An attractive hypothesis envisions that p.Met64Val and p.Met64Ile impair protein-protein interactions within the SRY-directed enhanceosome. Although components of this presumed assembly have not been defined, Met64 is exposed on one surface of the HMG box/DNA complex. Such exposure is not characteristic of other wedge residues (*i.e*., Trp98 within the core, intercalative side chain Ile68, and the latter’s aromatic partner Phe67 ^49^). In this model the WT side chain would play a dual role at the edge of the SRY-DNA interface and at a key protein-protein interface. A precedent for such overlapping surfaces has been provided by structures of SOX-POU-DNA ternary complexes ^84^. Testing this idea will require proteomic analysis of testis-specific enhanceosomes. Identification of SRY-interacting proteins and their functional validation have posed long-standing challenges ^14; 85^.

### Concluding remarks

We have described a novel inherited mutation in *SRY* at an invariant site in its conserved HMG box. This substitution alters the shape and volume of a “hydrophobic wedge” at a bent protein-DNA interface ^48^. The partial biological activity of p.Met64Val SRY (*ca*. 50% with respect to both *Sox9* enhancer occupancy and *Sox9* transcriptional activation) rationalizes both (a) the phenotype of the probands’ father and the normal prepubertal brother and (b) Swyer phenotypes of other affected 46, XY family members at the threshold of SRY function. Codon 64 in *SRY* represents a hot spot for clinical mutations leading to complete gonadal dysgenesis ^51^.

The majority of Swyer mutations in SRY represent *de novo* meiotic errors in paternal spermatogenesis leading to profound impairment of specific DNA binding or DNA bending^42^. Although inherited alleles are less common, such families are of interest for both clinical and mechanistic reasons. On the one hand, phenotypic variation (infertile 46, XY female, fertile male, or male requiring T supplementation) may represent either stochastic factors or modifier genes awaiting discovery. In addition, variants in other known genes or its regulatory non-coding regions involved in mammals sexual determination, should be ruled out. However, accordingly to this family pedigree, epigenetic events as well as digenic disorders might barely explain this intriguing phenomenon.

Besides, recognition of such pedigrees is important for genetic counseling and early assessment of family members at high risk for gonadal tumors—especially paradoxical secondary sexual characters may delay or confound genetic diagnosis. On the other hand, such “experiments of nature” provide rich opportunities for scientific discovery. In the present case it would be of particular interest to investigate structural and biophysical mechanisms leading to attenuated occupancies of SRY-responsive enhancers

## Acknowledgements

We thank Prof. M. Georgiadis (Indiana University) for helpful discussion regarding the structure of SRY. Y-S.C. was supported in part by a postdoctoral fellowship from the JDRF (New York, NY USA). M.A.W. is supported in part by the Robert A. Harris Chair in Biochemistry and Molecular Biology and Distinguished Professor Fund of Indiana University.

